# Evidence for multiple forms of heritable RNA silencing

**DOI:** 10.1101/2024.04.28.591487

**Authors:** Mary S. Chey, Pravrutha Raman, Farida Ettefa, Antony M. Jose

**Affiliations:** Department of Cell Biology and Molecular Genetics, University of Maryland, College Park, MD 20742

**Keywords:** epigenetics, transgenerational gene silencing, poly-UG RNAs, heredity, gene regulation, operon

## Abstract

Heritable gene silencing has been proposed to rely on DNA methylation, histone modifications, and/or non-coding RNAs in different organisms. Here we demonstrate that multiple RNA-mediated mechanisms with distinct and easily detectable molecular signatures can underlie heritable silencing of the same open-reading frame in the nematode *C. elegans*. Using two-gene operons, we reveal three cases of gene-selective silencing that provide support for the transmission of heritable epigenetic changes through different mechanisms of RNA silencing independent of changes in chromatin that would affect all genes of an operon equally. Different heritable epigenetic states of a gene were associated with distinct populations of stabilized mRNA fragments with untemplated poly-UG (pUG) tails, which are known intermediates of RNA silencing. These ‘pUG signatures’ provide a way to distinguish the multiple mechanisms that can drive heritable RNA silencing of a single gene.

## Introduction

Epigenetic control systems [1] that regulate the state of an organism evolve along with genome sequence. For a given genome sequence, numerous architectures formed by interacting regulators are compatible with heredity and evolution [2], although fewer may be compatible with particular functions. Nevertheless, detecting and analyzing alternative heritable states of regulatory architectures that can arise for the same genome sequence – i.e., heritable epigenetic changes – remains a challenge.

RNA silencing in the nematode *C. elegans* by double-stranded RNA (dsRNA) or by germline small RNAs called piRNAs can last for many generations without changes in genome sequence (reviewed in [3]). However, the susceptibility to this type of heritable epigenetic change and its transgenerational stability can vary dramatically depending on the target gene [4]. The maintenance of silencing for many generations is thought to require a positive feedback loop that includes the Argonaute protein HRDE-1 [5], which binds antisense small RNAs generated using RNA-dependent RNA polymerases (RdRPs) [6] and mRNA fragments stabilized through the addition of multiple UG dinucleotides (poly-UG or pUG RNAs) [7]. Since HRDE-1 can target nascent transcripts and recruit enzymes that can modify histones [5], a role for chromatin has also been proposed. Indeed, in many organisms, histone modification or DNA methylation is more widely studied in association with heritable epigenetic changes [8]. Here, we show that heritable epigenetic changes can persist even at single genes of operons independent of operon-level chromatin changes and that a diversity of mechanisms characterized by different molecular markers can underlie this heritable RNA silencing in *C. elegans*.

## Results and Discussion

Stable RNA silencing that is maintained for hundreds of generations can be induced by mating wild-type hermaphrodites without *mCherry* sequences to males with *mCherry* sequences expressed in the germline either as part of a transgene [4] or fused to the endogenous gene *sdg-1* [9]. However, the same *mCherry* sequence fused to the endogenous gene *mex-5* (*mCherry::mex-5* in Fig. 1*A*) remains expressed and is not susceptible to such mating-induced silencing [4]. To explore the mechanism(s) that promote transgenerational gene silencing, we used Mos1-mediated single-copy insertion (MosSCI) [10] to recreate a transgene that is susceptible to mating-induced silencing. This transgene *T* [4] encodes *mCherry::h2b* and *gfp::h2b* as two genes of an operon (Fig. 1*B*). Unlike the previously analyzed expressed version of *T* (one of 29% of such expressed isolates [4]), this newly inserted transgene did not show expression of either the mCherry::H2B or GFP::H2B upon insertion (designated *iTnew* in Fig. 1*B*; *i = ‘inactive’*). As expected [4], loss of the Argonaute protein HRDE-1 resulted in the re-expression of both proteins (Fig. 1*B*). Although similar insertion of another operon that has the location of *mCherry* swapped with that of *gfp* also failed to show any expression (designated *iTswap* in Fig. 1*C*), loss of HRDE-1 resulted in the recovery of mCherry::H2B fluorescence but not of GFP::H2B (designated as *giTswap* in Fig. 1*C*; *gi = ‘gfp inactive’*). Additional loss of proteins reported as required for heritable RNA silencing (the nucleotidyltransferase RDE-3/MUT-2, the Z-granule helicase ZNFX-1, or the P-granule associated protein DEPS-1) also did not result in re-expression of GFP::H2B. These observations reveal that while *Tnew* is silenced only by an HRDE-1-dependent mechanism (Fig. 1*D*, *left*), the two genes of *Tswap* are silenced by different mechanisms – one that is HRDE-1-dependent and another that is not (Fig. 1*D*, *right*). Although the underlying mechanism for this gene-selective silencing is unclear, chromatin-mediated mechanisms or pre-mRNA silencing that impact both genes equally can be excluded.

**Fig. 1.**
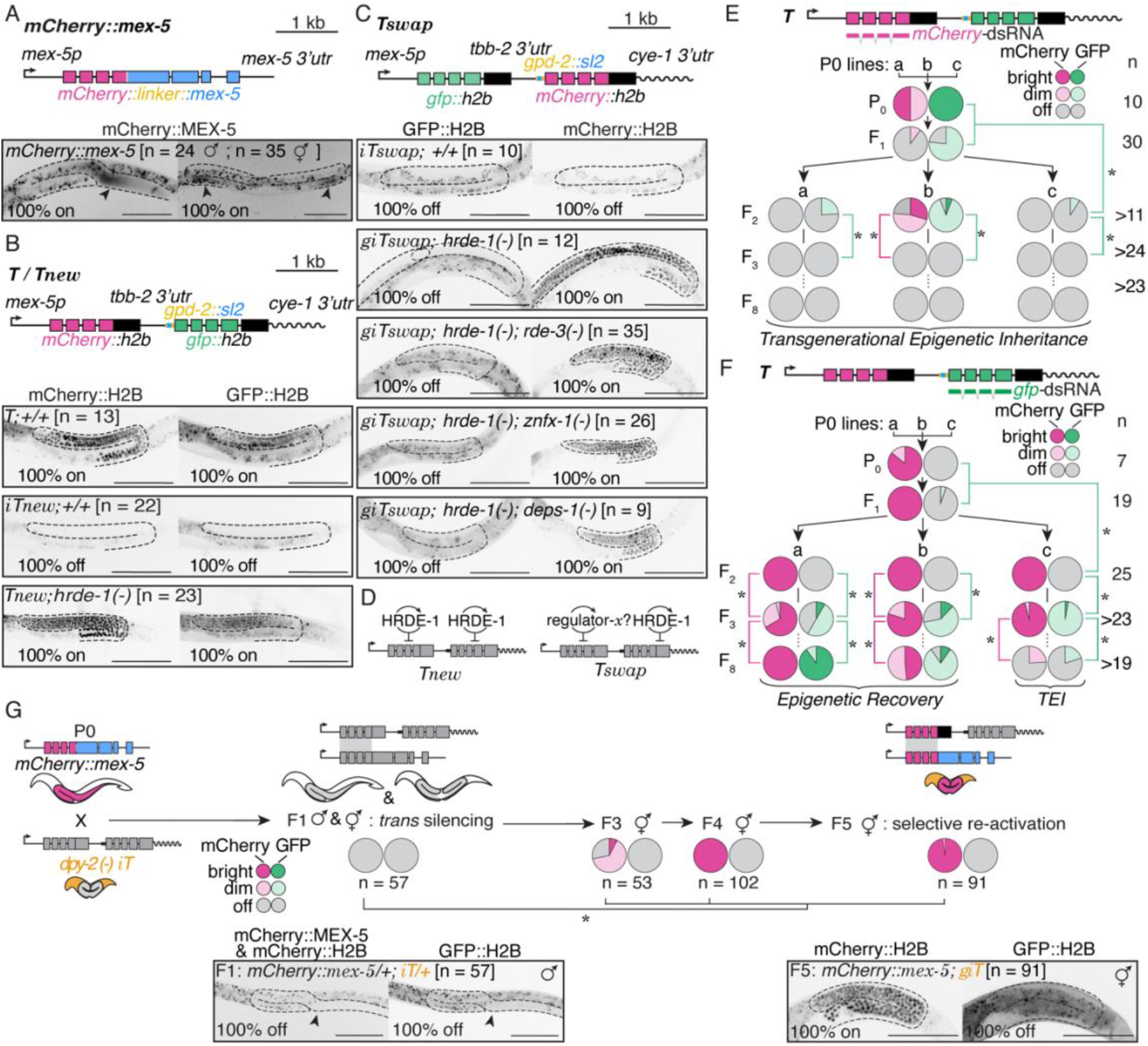
Gene-selective regulation of an operon can persist for multiple generations. (*A-C*) Expression from single-copy insertions of *mCherry* and/or *gfp* generated as a fusion with an endogenous gene (*mCherry::mex-5*) or as part of transgenes (*T, Tnew*, and *Tswap*). *Top*, Schematics. *Bottom*, Fluorescence (black) of mCherry and/or GFP fusion proteins within the germline (outlined) observed upon insertion of genes, if any, or recovered in various mutant backgrounds (premature stop codons in *hrde-1, rde-3/mut-2, znfx-1*, and/or *deps-1*). Scale bar = 100 μm, n indicates number of animals, arrowhead indicates site of mCherry::MEX-5 accumulation, and intestinal auto fluorescence can be seen as irregular back specks. (*D*) Summary of observations illustrating *hrde-1*-independent silencing of *gfp* but not *mCherry* in animals with *Tswap* versus the *hrde-1*-dependent silencing of both *gfp* and *mCherry* in animals with *T*. (*E* and *F*) Transgenerational dynamics of silencing in response to ingested dsRNA matching *mCherry* (*E*) or *gfp* (*F*). Fractions of animals showing silencing of each gene in each generation are indicated as pie charts. Lineages can show recovery of expression (epigenetic recovery) or long-term silencing (transgenerational epigenetic inheritance, TEI). Asterisks indicate *P* < 0.05 using the χ^2^ test when more than two categories are present or Wilson’s estimates for single proportions. (*G*) The expression of *mCherry::h2b* from *iT* generated by mating-induced silencing is selectively re-activated by *mCherry::mex-5* without re-activation of *gfp::h2b* despite initial *trans* silencing of *mCherry::mex-5*. Scale bars, arrowhead, n, and intestinal autofluorescence are as in (*A*); asterisks are as in (*E* and *F*). The DNA of *iT* was followed through crosses using a linked *dpy-2(-)* mutation (orange).

Similar gene-selective silencing was observed in two other cases: (1) upon ingestion of double-stranded RNA (dsRNA) and (2) upon exposure to expression of a homologous gene.

Silencing by dsRNA has revealed a mechanism for multigenerational gene silencing where the initial production of antisense small RNAs by RdRPs leads to histone modifications by methyltransferases ([11, 12] and reviewed in [3]). Consistently, when exposed to *mCherry*-dsRNA, animals with *T* showed selective silencing of *mCherry* in the exposed parents but silencing of both *gfp* and *mCherry* in descendants (Fig. 1*E*). These observations support the initial silencing of mRNA in the animals exposed to dsRNA followed by silencing of pre-mRNA and/or modification of chromatin in descendants (‘transgenerational epigenetic inheritance (TEI)’ in Fig. 1*E*). In contrast, exposure to *gfp*-dsRNA resulted in selective silencing of *gfp* in the exposed parents and in two additional generations before either recovery or silencing of both genes (‘epigenetic recovery’ or TEI in Fig. 1*F*). This selective silencing of *gfp* for three generations reveals an unstable form of heritable RNA silencing that is reminiscent of the 3-4 generations of silencing observed when the endogenous gene *oma-1* is targeted by injected *oma-1*-dsRNA [13].

The stable silencing of *T* initiated by mating is associated with the continuous production of antisense small RNAs that can silence genes of complementary sequence [4] – a phenomenon called *trans* silencing, which has also been observed using other transgenes (e.g., [14]). As expected, when *iT* is introduced into animals with *mCherry::mex-5*, the fluorescence from mCherry::MEX-5 is not detectable (Fig. 1*G*, F1 animals). However, in descendants with both *T* and *mCherry::mex-5*, fluorescence from mCherry::H2B, but not GFP::H2B, recovers (Fig. 1*G*, F5 animals). This selective re-activation of the *mCherry::h2b* gene in *T* also reveals a mechanism for silencing *gfp::h2b* mRNA without the joint loss of both genes as would be expected for silencing at the level of pre-mRNA or chromatin.

Heritable RNA silencing is associated with the production of pUG RNAs, which are mRNA fragments stabilized by the addition of multiple UG dinucleotides [7]. We used an RT-PCR-based assay [7] where a gene-specific 5’
s primer and pUG-specific 3’ primer are used to amplify cDNA matching the population of pUG RNA sequences made by cleaving mRNA downstream of the start codon (Fig. 2*A*). Strikingly, we found different populations of sequences were amplified using primers for the *mCherry* gene from each strain (Fig. 2*B*, *left*). Similarly strain-specific patterns were obtained using primers for *gfp* (Fig. 2*B*, *middle*), although the patterns were distinct from those observed for *mCherry*. These patterns of pUG RNAs were characteristic of each strain and were largely reproducible when the total RNAs used were prepared from two different populations (Fig. 2*C*). The presence of detectable pUG RNAs in total RNA from animals that show expression within the germline (*gfp* and *mCherry* from *T* and *mCherry* from *giTswap; hrde-1(-)* in Fig. 2*B* and 2*C*) suggests that either pUG RNAs are made within the germline but are insufficient for silencing or that these pUG RNAs are made in a somatic tissue where the *gfp* and *mCherry* sequences are silenced. Therefore, these distinct pUG signatures are molecular indicators of different states of stable expression or silencing for the same open-reading frame. Given the recent discovery that the cleavage of mRNA for pUG RNA production during RNAi of a somatic target gene is restricted to regions that match the dsRNA sequence called pUG zones [15], the different pUG signatures observed could be the result of distinct pUG zones created by different primary triggers of pUG RNA production. However, the underlying regulatory architectures that cause the observed molecular differences are currently unknown.

**Fig. 2.**
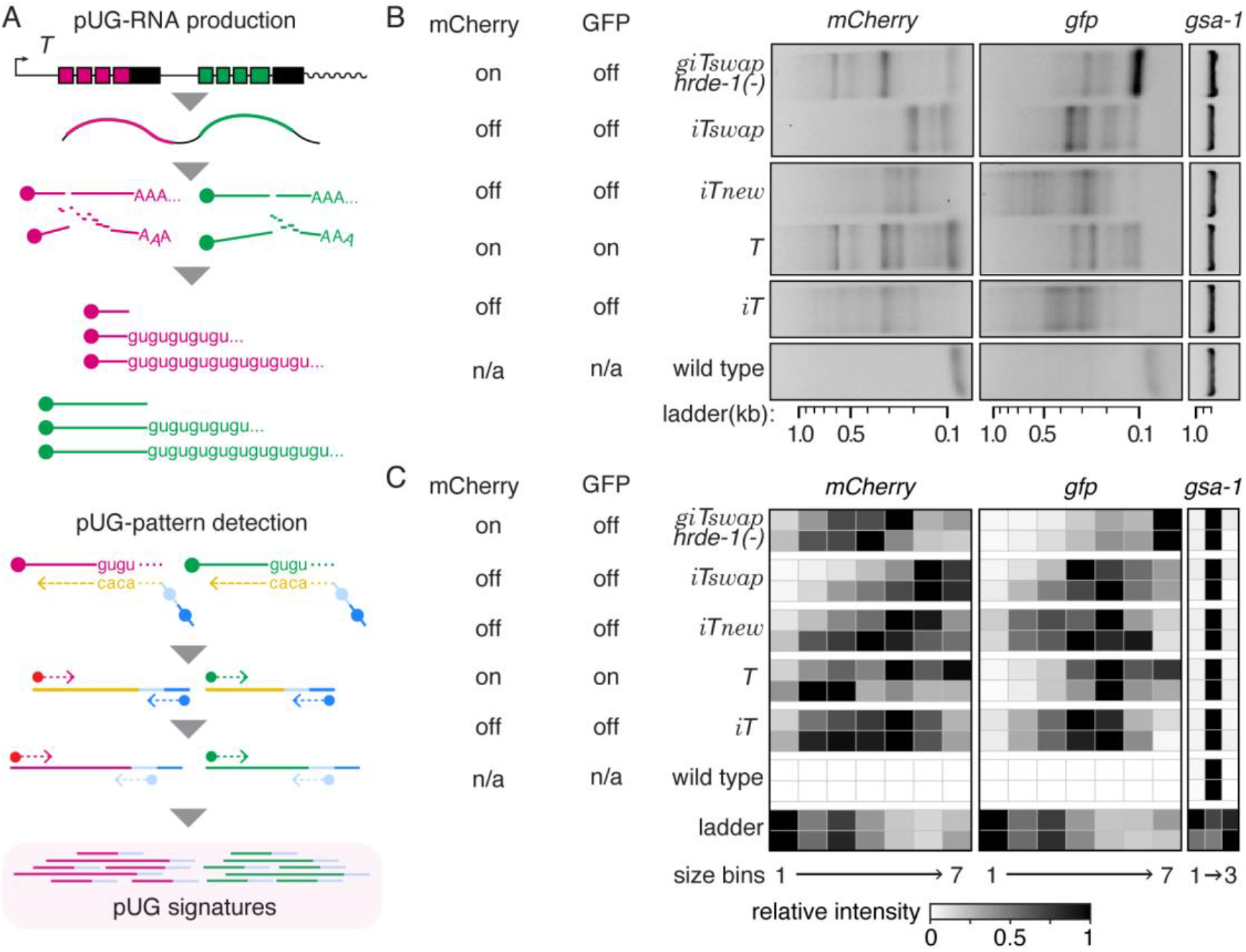
Heritable RNA silencing of the same open-reading frame can generate distinct populations of pUG RNA. (*A*) Schematic of pUG RNA production from *T* (*top*) and the detection of pUG signatures using RT followed by nested PCR (*bottom*). (*B*) Patterns of pUG RNAs derived from *mCherry* (*left*) and *gfp* (*middle*) sequences in animals with single-copy transgenes (*iT, iTnew, T, iTswap*, and *giTswap; hrde-1(-)*) or without (wild type). The poly-UG region of *gsa-1* mRNA serves as a positive control (*right*). (*C*) Heatmap of pUG RNA patterns detected in total RNA isolated from biological replicates of strains in (*B*) created by summing the intensities of amplified DNA from 0.1 to 1 kb into 7 size bins and assigning each bin a relative intensity per lane. See Supporting Information for details.

Since each regulatory architecture can be driven by different epigenetic changes into many alternative states [2], their unambiguous analysis requires keeping track of both genetic and epigenetic states. For example, since the genotypes of *T, iT*, and *iTnew* are all the same, they can be designated similarly using established nomenclature [16] (say, *labSi#* where ‘Si’ denotes MosSCI insertion). In contrast, their different epigenetic states need different designations (i.e., {*Epi-labSi#(e-lab1)*}, {*Epi-labSi#(e-lab2)*}, and {*Epi-labSi#(e-lab3)*}, respectively; see Strains in Supporting Information for examples and [17] for a discussion). Analyzing the diversity of heritable epigenetic changes that can arise at single genes could reveal the logic of heritable epigenetic effects and enable the design of regulatory circuits with predictable heredity.

## Materials and Methods

DNA insertions were generated using MosSCI or Cas9-mediated genome editing. Mutations in known regulators of heritable RNA silencing were introduced through Cas9-mediated genome editing or genetic crosses. Bacteria expressing *gfp*-dsRNA or *mCherry*-dsRNA were used to examine transgenerational dynamics in response to feeding RNAi. Fluorescence from mCherry or GFP fusion proteins in various strains was captured using a Nikon AZ100 microscope and images were identically adjusted using FIJI (NIH). Patterns of pUG RNAs present in total RNA were measured using reverse transcription followed by nested PCR and separating the populations of amplified DNA on a 1% agarose gel. See Supporting Information for detailed materials and methods.

## Acknowledgements

We thank Rui Yin for making the plasmid with *Tswap* DNA; Tom Kocher, Bill Snell, and members of the Jose Lab for comments on the manuscript; and the *Caenorhabditis elegans* Genetic Stock Center for some worm strains. This work was supported in part by National Institutes of Health Grants R01GM111457 and R01GM124356, and National Science Foundation Grant 2120895 to A.M.J.

