## Supplementary Information for "Evidence for multiple forms of heritable RNA silencing"

**Extended Materials and Methods**

**Strains.** Different stable phenotypes associated with the same genotype are designated as epigenetic states using a nomenclature that includes a brief description of history as proposed earlier [1].

| **name** | **shorthand (if any)** | **genotype {*Epi-locus(e-allele [history])*}** |
|---|---|---|
| N2 | +/+ | wild type |
| GE1708 | | *dpy-2(e8) unc-4(e120) II* |
| EG4322 | | *ttTi5605 II; unc-119(ed9) III* |
| EG6787 | *T* | *oxSi487 [mex-5p::mCherry::h2b::tbb-2 3'utr::gpd-2 operon::sl2::gfp:: h2b::cye-1 3'utr + unc-119(+)] II {Epi-oxSi487(e-jam1 [both mCherry::H2B and GFP::H2B were expressed upon insertion of oxSi487 using MosSCI])}* |
| AMJ552 | *iT* | *oxSi487 dpy-2(jam33[*e8]) {Epi-oxSi487(e-jam2 [both mCherry::H2B and GFP::H2B were silenced by mating males with expressed copies of oxSi487 to wild-type hermaphrodites without the transgene and then re-homozygosing the silenced transgene])}* |
| AMJ1652 | *iTnew* | *jamSi83 [mex-5p::mCherry::h2b::tbb-2 3'utr::gpd-2 operon::sl2::gfp:: h2b::cye-1 3'utr + unc-119(+)] II {Epi-jamSi83(e-jam3 [both mCherry ::H2B and GFP::H2B were silenced upon insertion of jamSi83 using MosSCI])}* |
| AMJ581 | *T dpy-2(-)* | *oxSi487 dpy-2(e8)* II*; unc-119(ed3)? III {Epi-oxSi487(e-jam1 [expression of both mCherry::H2B and GFP::H2B from oxSi487 was preserved by mating hermaphrodites with the transgene to males with dpy-2(e8) unc-4(e120)/++ males and selecting oxSi487 dpy-2(e8) animals])}* |
| AMJ1665 | *Tnew; hrde-1(-)* | *jamSi83; hrde-1(jam303) {Epi-jamSi83(e-jam4 [both mCherry::H2B and GFP::H2B were re-expressed upon introducing a premature stop codon into hrde-1 in AMJ1652])}* |
| AMJ844 | *iT dpy-2(-)* | *oxSi487 dpy-2(e8) {Epi-oxSi487(e-jam2)}* |
| AMJ1208 | *mCherry::mex-5* | *mex-5(jam197[Pmex-5::mCherry::mex-5::mex-5 3'utr])* |
| AMJ1629 | *iTswap* | *jamSi79 [mex-5p::gfp::h2b::tbb-2utr::mCherry::h2b::cye-1utr]* II *{Epi-jamSi79(e-jam5 [both GFP::H2B and mCherry::H2B were silenced upon insertion of jamSi79 using MosSCI])}* |
| AMJ1793 | *giTswap; hrde-1(-)* | *jamSi79; hrde-1(jam342) {Epi-jamSi79(e-jam6 [mCherry::H2B, but not GFP::H2B, was selectively re-expressed upon introducing a premature stop codon into hrde-1 in AMJ1629])}* |
| AMJ1794 | *giTswap; hrde-1(-); rde-3(-)* | *jamSi79; hrde-1(jam342); rde-3(jam343) {Epi-jamSi79(e-jam7 [mCherry::H2B, but not GFP::H2B, was selectively re-expressed upon introducing a premature stop codon into rde-3 in AMJ1793])}* |
| AMJ1795 | *giTswap; hrde-1(-); znfx-1(-)* | *jamSi79; hrde-1(jam342); znfx-1(jam344) {Epi-jamSi79(e-jam8 [mCherry::H2B, but not GFP::H2B, was selectively re-expressed upon introducing a premature stop codon into znfx-1 in AMJ1793])}* |
| AMJ1796 | *giTswap; hrde-1(-); deps-1(-)* | *jamSi79; hrde-1(jam342); deps-1(jam345) {Epi-jamSi79(e-jam9 [mCherry::H2B, but not GFP::H2B, was selectively re-expressed upon introducing a premature stop codon into deps-1 in AMJ1793])}* |
| AMJ1421 | *mCherry::mex-5; giT dpy-2(-)* | *oxSi487 dpy-2(e8); mex-5(jam197) {Epi-oxSi487(e-jam10 [expression of mCherry::H2B, but not GFP::H2B, was selectively re-activated upon multi-generational exposure to mex-5(jam197))}* |

**Oligonucleotides.** Primers used for RT or PCR, crRNAs and homology repair templates used for Cas9-mdiated genome editing are indicated below.

| name | sequence | use |
|---|---|---|
| P1 | ataaggagttccacgcccag | genotyping of operon variants [*oxSi487*, *jamSi83*, and *jamSi79*] |
| P2 | ctagtgagtcgtattataagtg | |
| P3 | tgaagacgacgagccacttg | |
| P4 | gtgtcgaagttgtctcgcag | genotyping *hrde-1(jam342)* |
| P5 | acggtatccaccacgagag | |
| P6 | cgctaacacaactcctttgc | genotyping *rde-3(jam343)* |
| P7 | ccacgttggattgttcttcg | |
| P8 | tccgttgacagaggttacatgc | genotyping *deps-1(jam345)* |
| P9 | agcgtcttccagcagaaatg | |
| P10 | cacaacggctcgatacaagg | genotyping for *znfx-1(jam344)* |
| P11 | cgttctgcccgtttgatcc | |
| P12 | gctatggctgttctcatggcggcgtcgccatattctacttcacacacacacacaca | Reverse Transcription primer for detecting pUG signatures |
| P13 | gctatggctgttctcatggc | reverse adapter 1 for detecting pUG signatures |
| P14 | ggcgtcgccatattctactt | reverse adapter 2 for detecting pUG signatures |
| P15 | gagttctacgatcacattct | Primers for amplification of *gsa-1* controls when looking for pUG signatures |
| P16 | cacttgctggaaagacaagg | |
| P17 | atggtctccaagggagag | Primers for detecting pUG signatures from mCherry |
| P18 | gagaggaggataacatggct | |
| P19 | atgagtaaaggagaagaacttttca | Primers for detecting pUG signatures from gfp |
| P20 | ttcactggagttgtccca | |
| P21 | gaaaguuucaacgcguuuua | crRNA for introducing premature stop into *rde-3* |
| P22 | cagacguuuggcuauacgcc | crRNA for introducing premature stop into *deps-1* |
| P23 | aaugaggauuacuuacaauu | crRNA for introducing premature stop into *hrde-1* |
| P24 | cauuauuauaggcgccgucu | crRNA for introducing premature stop into *znfx-1* |
| P25 | cgtttcacaaatttaacggaaagtttcaatgagttttaagggatcacgaggacgatttca | HRT for introducing premature stop into *rde-3* |
| P26 | gcgacccgtgcggagccagacgtttgactatacgcctggattcgattcgaaactaccatg | HRT for introducing premature stop into *deps-1* |
| P27 | aaagctcaacaatgaggattactta**t**aatttggtaatgccacattcccacgtctttctcc | HRT for introducing premature stop into *hrde-1* |
| P28 | gcgacccgtgcggagccagacgtttgactatacgcctggattcgattcgaaactaccatg | HRT for introducing premature stop into *znfx-1* |
| P29 | ggcagaatgtgaacaagactcg | Genotyping transgenes after integration using MosSCI |
| P30 | gctgcagccctcttaatgc | |
| P31 | gaagggcgccctaactttg | |

**Plasmids.** For making *gfp*-dsRNA ([2] gift from the Hamza lab (University of Maryland, School of Medicine) and verified using Sanger sequencing by Ed Traver, Jose lab) or *mCherry*-dsRNA, *gfp* and *mCherry* cDNA sequences were cloned into the pL4440 vector backbone, which has flanking T7 promoters, and transformed into HT115 *E. coli*. The *mCherry*-dsRNA plasmid was created by amplifying the *mCherry* exon sequences from EG6787 and then using Gibson Assembly (NEBuilder HiFi DNA Assembly Master Mix, New England Biolabs (NEB)). The resulting plasmid was verified using Sanger sequencing.

For homology repair after Mos1 excision, the operon sequences were amplified from pCFJ359 (containing the sequences of *T* [3]) and pSD5 (containing the sequences of *Tcherry* [4]) using Phusion High Fidelity polymerase (NEB, catalog no. M0530S) and cloned using Gibson Assembly followed by transformation into DH5$\alpha$ *E. coli*. Single colonies were isolated, and sequences were verified using Sanger sequencing.

**Feeding RNAi.** Animals were exposed to dsRNA targeting either *gfp* or *mCherry* only in the P0 generation as described for *gfp*-dsRNA in Devanapally et al. [4]. Briefly, worms were grown at 20°C on nematode growth media (NGM) plates supplemented with 1 mM IPTG (Omega Bio-Tek) and 25 µg/ml Carbenicillin (MP Biochemicals). Age-matched P0 animals were fed control RNAi (HT115 bacteria with pL4440), *mCherry*-dsRNA or *gfp*-dsRNA for 24 hrs post L4, and controls were scored alongside. P0 animals and their untreated descendants growing on OP50 were analyzed for mCherry::H2B and GFP::H2B expression from *T*. L4-staged animals were scored in each generation by imaging (except F2 animals after *gfp* RNAi, which was scored by eye). Unimaged L4 siblings were passaged without bias in each generation to obtain progeny for the next generation.

**Strain creation.** Genetic crosses, Mos1-mediated single copy insertion (MosSCI) [5], or Cas9-mediated genome editing were used for creating various strains. In cases where a locus was susceptible to mating-induced silencing, care was taken to preserve the susceptible transgene in hermaphrodites to avoid such silencing [4].

*Genetic crosses.* L4-staged males (~9 animals) containing *mCherry::mex-5* were crossed with *iT dpy-2(-)* hermaphrodites (3 animals). F1 L4-staged nonDpy hermaphrodites and males were isolated and either imaged as L4-staged or 24-48 hours post L4 animals. Isolated F1 nonDpy hermaphrodites were also allowed to lay self-progeny. F2 hermaphrodites were singled out, allowed to lay progeny, and genotyped after ~3.5 days to identify *mCherry::mex-5; iT dpy-2(-)* homozygotes. F3 hermaphrodites were similarly passaged and genotyped to confirm homozygosity. Subsequent generations were obtained through unbiased passaging of animals and unpassaged siblings were imaged in each generation.

*MosSCI.* EG4322 adult animals were injected 24 hours after the L4 stage with a 10 µl mix containing pCFJ601 (50 ng/µl), pMA122 (10 ng/µl), and a plasmid-containing operon of interest (55-60 ng/µl). Injected animals were isolated one to a plate and allowed to have progeny. After ~10 days, when the plate was crowded with starved animals, they were screened for nonUnc animals. The nonUnc animals were subject to heat shock in a water bath at 34ºC for 2.5 hrs. After 2-3 days of recovery at 20ºC, remaining nonUnc animals, if any, were isolated onto NGM plates and allowed to have progeny. Each parent nonUnc hermaphrodite was lysed and the lysates were split three ways to genotype for the left junction (P29 and P31) and right junction (P2 and P30) of integration, as well as for homozygosity (P29 and P30). Genomic DNA was prepared from plates of animals with successful MosSCI integration, and different primer sets were used to amplify the entire integrated sequence by PCR for Sanger sequencing. Sequencing of the operon inserts revealed no mutations anywhere except a mis-sense mutation within the *h2b* sequences (*his-58*: c.C314T **|** p.A105V) fused with *gfp* in AMJ1629. The relevance of this mutation for the observations on *Tswap*, if any, is unknown.

*Cas9-mediated genome editing.* Adult animals were injected 24 hours after the L4 stage with 10 µl mix containing tracrRNA (~9 pmol/µl), crRNA for the gene of interest (~10-47 pmol/µl), *dpy-10* crRNA (~30 pmol/µl) or pRF4 (40 ng/µl), and Cas9 protein (1.6 pmol/µl; PNA Bio catalog no. CP01). Homology templates for generating *dpy-10(-)* were single-stranded DNA oligos [4]. However, homology templates to generate *mCherry::mex-5* were amplified from plasmids containing the reporter sequence using Phusion High Fidelity polymerase (New England Biolabs catalog no. M0530S) and gene-specific primers with ~35 bp of overhang homologous to the site of integration. PCR products were purified using NucleoSpin Gel and PCR Clean-up (Macherey-Nagel, catalog no. 740609.250).

**Imaging.** Animals of the L4 stage or between 24-72 hours after the L4 stage were mounted on a slide after paralyzing them using 3 mM levamisole (Sigma-Aldrich, Cat# 196142), imaged under non-saturating conditions (Nikon AZ100 microscope and Photometrics Cool SNAP HQ2 or Prime BSI Express camera), and categorized into a maximum of three groups (bright, dim and off). A C-HGFI Intensilight Hg Illuminator was used to excite GFP (filter cube: 450 to 490 nm excitation, 495 nm dichroic, and 500 to 550 nm emission) or mCherry (filter cube: 530 to 560 nm excitation, 570 nm dichroic, and 590 to 650 nm emission). Sections of the gonad that were not obscured by autofluorescence from the intestine were examined to classify GFP and mCherry fluorescence. The expression of mCherry::MEX-5 after mating-induced silencing or *trans* silencing was categorized as 'on' only if fluorescence was visible when the LUTs were adjusted to a range of 0-5,000. Intestinal autofluorescence was more appreciable when imaging GFP than when imaging mCherry. In the case where fluorescence was scored by eye (F2 animals after *gfp*-dsRNA feeding), it was done at fixed magnification and zoom using an Olympus MVX10 microscope for GFP (filter cube: 460 to 480 nm excitation, 485 nm dichroic, and 495 to 540 nm emission) and mCherry (filter cube: 535 to 555 nm excitation, 565 nm dichroic, and 570 to 625 nm emission). Representative worm images were adjusted to different brightness/contrast to view different germline gene expression patterns using FIJI (NIH). All images being compared were identically adjusted for display.

**pUG signatures.** DNA amplicons corresponding to pUG RNA populations were amplified essentially as described in Shukla et al. [6] with the additional analysis of final patterns of the amplified populations using FIJI (NIH, version 2.4.0). All worm strains were grown on 35 mm NGM plates at 20°C seeded with 100 µl of *E. coli* OP50. Just before all the *E. coli* was consumed, ~6 plates of worms were washed three times with M9 buffer and pooled using centrifugation at 14,000 rpm for 30 sec after each wash, yielding a ~100 µl pellet of worms. The worm pellet was then mixed with 1 ml of TRIzol (Fisher Scientific) and subject to 3 freeze/thaw cycles in liquid nitrogen. RNA was extracted according to manufacturer's protocol using chloroform and precipitated along with 1-2 µl of glycogen (5 µg/µl in Ambion cat. #AM9510 or 20 µg/µl in Invitrogen cat. #10814-010). Reverse transcription was performed using Superscript III (Invitrogen) and RT primer P12 using ~2000 ng of total RNA. Subsequent nested PCR was performed using ~2 µl of cDNA with the first round of amplification done using Phusion Taq Polymerase (NEB) in a final reaction volume of 20 µl for 25 cycles. The first set of gene-specific forward primers were used for each target gene (*mCherry* – P17, *gfp* – P19, and *gsa-1* – P15) with the adapter 1 reverse primer (P13). The PCR product from round 1 was diluted 100-fold and 1 µl of the diluted product was used for the second round of PCR in a total reaction volume of 50 µl. The second set of gene-specific forward primers (*mCherry* – P18, *gfp* – P20, and *gsa-1* – P16) were used with adapter 2 reverse primer (P14). The PCR products (~20 µl for *mCherry* and *gfp*, and ~8 µl for *gsa-1*) were combined with 6X loading dye, run on a 1% agarose gel and imaged using GelDoc (BioRad). All gels included an N2 control from which only a *gsa-1* amplicon is expected. A screenshot was used to generate a .png file of each gel before quantifying pUG signatures. The intensities of all bands were measured using a line of similar length through the middle of each lane using the 'Plot Profile' tool in FIJI (NIH) and processed using a custom script (R, version 4.2.2) to obtain a final heatmap (Fig. 2*C*). For the heatmap, the intensities of the N2 lane were subtracted from that at the corresponding positions of each sample lane (background subtraction). These background-subtracted intensities were then combined into 7 bins ranging from above the largest band of a 100-bp DNA ladder (NEB). The specific bins for all gels in arbitrary units of distance are as follows: one = 1 to 70, two = 71 to 100, three = 101 to 150, four = 151 to 180, five = 181 to 220, six = 221 to 260, and seven = 261 to 'bottom'. The 'bottom', which is just below ~100 bp, varied slightly for each pair of gels: 309 for *gfp*, 294 for *mCherry* and 296 for *gsa-1*. These binned intensities were then normalized for each lane by dividing each intensity by the maximal for that lane. With these normalized data, the heatmap was generated by setting maximum to black and minimum to white for each lane. While intensities cannot be compared across different genes because of differences in PCR efficiency, characteristic patterns for the same gene in a strain that remain comparable across biological replicates reveal the pUG signature for that gene in that strain.

**Statistics.** When two categories (on or off) were present in both datasets being compared, Wilson's estimates for single proportions was used and when more than two categories were present (on, off, and dim), the $\chi 2$ test was used [4].
